# Single Cell Transcriptional Profiling of Phox2b-Expressing Geniculate Ganglion Neurons

**DOI:** 10.1101/812578

**Authors:** Catherine B. Anderson, Eric D. Larson

**Author notes:** Author Roles: CBA performed experiments and analyzed data. EDL conceived and executed the study, performed experiments, performed data analyses, and wrote the manuscript. All authors reviewed and approved the final manuscript. Corresponding Author: Eric D. Larson, University of Colorado, Department of Otolaryngology, 12700 E. 19^th^ Ave, MS 8606, Aurora, CO 80045.

## Abstract

The sense of taste is fundamental for survival as harmful substances can be discriminated and prevented from entering the body. Taste buds act as chemosensory sentinels and detect bitter, salty, sweet, sour, and umami substances and transmit signals to afferent nerve fibers. Whether a single gustatory nerve fiber selectively is responsive to a single taste modality (through taste receptor cell activation) is a point of contention in the field.. In the present study, we present a method for single cell RNA sequencing of gustatory geniculate ganglion neurons and compare the results obtained to two prior published works. Additionally, independent reanalysis of the raw data from these previous studies confirms molecular heterogeneity of ganglion neurons. Multiple gustatory clusters are found, and we compare cluster markers identified by the original works and those identified in the present study. Across all datasets and analyses, specific clusters show a high degree of correlation including a somatosensory cluster (*Phox2b*-, *Piezo2*+, *Fxyd2*+), a potential sweet-best cluster (*Phox2b*+, *Spon1*+, *Olfm3*+), and a potential sour-best cluster (*Phox2b*+, *Penk*+, *Htr3a*+). Additionally, a putative mechanosensitive gustatory cluster with an unknown functional role is identified (*Phox2b*+, *Piezo2*+, *Calb1*+). Other gustatory clusters (*Phox2b*+) are more varied across analyses, but are marked by *Olfm3*. Which, if any, clusters comprise umami-best, bitter-best, or salty-best fibers will require further study.

## Introduction

Taste buds detect sapid molecules in the mouth and are the initiators of gustation. Each taste bud contains 50-100 mature taste cells, which are classified as three separate types. Type I cells are the least understood, but act as supporting cells and may transduce some portion of salty taste. Type II cells are well-understood and transduce signals in response to bitter, sweet, or umami stimuli. Type II cells communicate with afferent nerve fibers through release of ATP via non-traditional synaptic mechanisms (Finger et al., 2005; Huang et al., 2007; Romanov et al., 2007, 2018; Taruno et al., 2013). Type III cells form classical synapses with afferent nerve fibers and are responsible for sour and portions of salt taste detection. Due to the different nature of how Type II and Type III cells communicate with afferent fibers, it is likely that the innervating fibers have different molecular and physiological properties. Indeed, we have shown that afferent fibers that express the serotonin receptor 5-HT3A preferentially contact serotonergic Type III cells (Stratford et al., 2017). This subset of afferent fibers responds to exogenous serotonin as well as ATP while other gustatory fibers respond only to ATP (Larson et al., 2015). These data suggest at least 2 subpopulations of gustatory afferent nerve fibers.

We previously showed that the majority of geniculate ganglion neurons show excitatory responses to exogenously applied ATP and about 25% to serotonin via P2X2/P2X3 and 5HT3A receptors, respectively (Larson et al., 2015; Vandenbeuch et al., 2015). Older studies using patch clamp electrophysiology of rat gustatory geniculate ganglion neurons also showed excitatory responses to ACh, serotonin, substance P, and GABA in a small percentage of cells (King and Bradley, 2000; Koga and Bradley, 2000). These data suggest that multiple classes of neurons exist at the physiological level and that likely, these classes could be reflected at the molecular level.

Lingual taste fields are innervated by branches of the facial (cranial nerve VII) and glossopharyngeal nerve (cranial nerve IX). The anterior tongue receives input from the chorda tympani branch of the facial nerve. The cell bodies of the chorda tympani reside in the geniculate ganglion. In addition to the chorda tympani, this ganglion houses the cell bodies of the greater superficial petrosal and the nervus intermedius which receive gustatory input from the soft palate and somatosensory input from the external auditory meatus (Mtui et al., 2011).

Gustatory geniculate ganglion neurons are delineated by expression of the transcription factor, *Phox2b* (Dvoryanchikov et al., 2017; Ohman-Gault et al., 2017; Rios-Pilier and Krimm, 2019) which molecularly defines a neuron as gustatory or non-gustatory. The first study to characterize the transcriptome of single geniculate ganglion neurons utilized the Fluidigm C1 to profile 96 neurons from wildtype mice (Dvoryanchikov et al., 2017). They found about 2/3 of the cells express *Phox2b*, consistent with histological studies. Within the gustatory cells, three main clusters were identified, all of which expressed *Phox2b* and *P2rx3*. Other cluster markers included T1: *Olfm3*, *Itm2a*, *Hspb3*; T2: *Cd302*, *Kcnip1*, *Slc39a11*; T3: *Lypd1*, *Penk*, and *Trhr*. The authors then characterized the physiological response profiles of isolated ganglion neurons to exogenous ATP and 5-HT to match functional profiling with transcriptional profiling and demonstrated multiple physiological profiles that nicely matched expression of P2X and 5-HT receptor mRNA. This study was the first to characterize geniculate ganglion neurons at the molecular level using scRNA-seq.

More recently, Zhang et al. performed scRNA-sequence analysis on a larger cohort of geniculate ganglion neurons (Zhang et al., 2019). They profiled over 400 neurons using the Cel-seq (Hashimshony et al., 2012) method and identified 7 clusters. Based on cluster markers, they made multiple transgenic and knockout mice to test the role of certain transcripts in taste detection. Molecular and physiological evidence is presented for classes of sweet-best and sour-best neurons, marked predominantly by expression of *Spon1* and *Penk* transcripts, respectively. Using a floxed Gcamp model, the authors recorded in vivo calcium signals in geniculate ganglion neurons and showed that *Spon1* and *Penk* neurons selectively responded to sweet and sour lingual stimuli, respectively. Knockout of *Cdh4* and *Cdh13* eliminated detection of umami and bitter molecules, respectively, as assessed by two-bottle taste preference testing. Inhibition of *Egr2*-neurons using a *Egr2*-cre/Tetanus toxin model eliminated detection of NaCl. However, it is unclear how protein products of these transcripts are involved in taste signaling.

In the present study, we performed transcriptional profiling of geniculate ganglion neurons using an alternative method. We show that the neurons we collected expressed many transcripts overlapping with currently published datasets. Additionally, we performed a secondary analysis of published datasets from Dvoryanchikov et al 2017 and Zhang et al 2019 to further explore geniculate ganglion neuron clusters in attempt to understand more about the role of ganglion neurons in taste perception. Lastly, we show that combining all three datasets preserves cell clustering using the new analysis methods in Seurat V3 (Butler et al., 2018; Stuart et al., 2019).

## Materials and Methods

### Animals

All animal procedures were conducted under a protocol approved by the University of Colorado Animal Care and Use Committee. Phox2bcre mice were purchased from Jax (B6(Cg)-Tg(Phox2b-cre)3Jke/J; stock # 016223) and bred to Rosa26/tdTomato mice (B6.Cg-Gt(ROSA)26Sortm14(CAG-tdTomato)Hze/J; stock # 007914). Crossed mice of both gender were used between the ages of 8 and 24 weeks.

### Immunofluorescence

Animals were anesthetized using urethane and transcardially perfused with 4% paraformaldehyde in phosphate buffer (0.1M; PB) or euthanized with carbon dioxide. Geniculate ganglia and tongues were extracted and fixed for 1 or 4 hours at room temperature before incubating in 20% sucrose in PB overnight at 4°C. Tissue was embedded and frozen in Optimal Cutting Temperature (OCT) and sectioned at a thickness of 12-16 μm using a cryostat. After > 24 hours at −20°C, slides were thawed, rehydrated with PBS, and blocked using 2% normal donkey serum in blocking buffer (PBS plus 0.3% TritonX100, 1% BSA). Primary antibodies were applied overnight at 4°C. After thorough washing fluorescent secondary antibodies were applied at room temperature for 3 hours before counterstaining with DAPI and mounting with FluoroMount G (Southern Biotech). Slides were imaged using a Leica SP8 using 20x (NA 0.75) and 63x (NA 1.4) oil-immersion objectives.

### Single cell preparation

Geniculate ganglia of three carbon dioxide-euthanized mice were rapidly extracted and placed in minimum essential medium with Earle’s balanced salts (MEM/EBSS; Hyclone) supplemented with 1.25 mg/ml trypsin (Sigma-Aldrich) and 2.5 mg/ml collagenase A (Roche Diagnostics) for 30 minutes at 35°C. Digested ganglia were washed with MEM/EBSS, and gently triturated with a fire-polished glass pipette. The suspension was passed through a 50 μm filter and loaded as the top layer of a Percoll gradient (12.5 and 28%, Sigma-Aldrich; Malin et al., 2007). The gradient was spun at 1300xg for 10 minutes using a swinging bucket rotor. The cell pellet at the bottom of the tube was washed with MEM/EBSS before resuspending in 2 mL MEM/EBSS.

### Single cell capture and RNA extraction

A QiaScout Microraft Array (Qiagen) was coated with poly-D-lysine (0.02 mg/ml) for 2 hours and washed with MEM/EBSS. The geniculate ganglion cell suspension was placed in the microraft array and the cells allowed to settle for 20 minutes. The microraft array was secured using a custom set up to the stage of an Olympus IX71 inverted microscope equipped with a 10X objective and the QiaScout instrument (Qiagen). Cells were visually inspected and microrafts containing a single tdTomato-positive (or negative in some cases) geniculate ganglion neuron were extracted and transferred to a PCR tube containing lysis buffer as part of the QIAseq FX Single Cell RNA Library Kit (Qiagen). Once in lysis buffer, cells were lysed by heating the tube to 95°C for 3 minutes followed by an infinite hold at 4°C until 24 individual samples were collected. The collection protocol was performed across two independent experiments, resulting in 48 individual sequencing libraries (42 tdTomato+, 6 tdTomato-).

### Library preparation and sequencing

Sequencing libraries were prepared using the QIAseq FX Single Cell RNA Library Kit (Qiagen) according to manufacturer instructions. Cells were lysed, followed by gDNA removal, reverse transcription, ligation, and whole transcriptome amplification. After amplification, libraries were enzymatically fragmented (~300 bp fragments) followed by adapter ligation. Library size was evaluated using a DNA tapestation and concentration was determined using qPCR (QIAseq Library Quant Array). Libraries were pooled at equimolar concentration and sequenced using an Illumina MiSEQ to generate 2×151 bp reads.

### Sequencing analysis

Quality and adapter trimming was performed using BBDuk (Bushnell). Salmon (v0.14.0) was used to quantify transcript expression using trimmed reads and the Ensembl GRCm38.p6 (version 92) transcriptome (Patro et al., 2017; Zerbino et al., 2018). Transcript expression was summarized at the gene level and input to R using TxImport (Soneson et al., 2016; R Core Team, 2019). A custom R script was used to visualize gene expression in the dataset. Samples were kept if expression of *Snap25* or *Uclh1* was > 5 TPM. Discarded samples were further interrogated for expression of neuron-specific genes, but they were rarely present.

### Sequencing analysis of published datasets

Single cell sequencing FASTQ files were obtained from NCBI Gene Expression Omnibus (GSE102443 (Dvoryanchikov et al., 2017) and GSE135801 (Zhang et al., 2019)). Raw reads from Dvoryanchikov et al. were filtered, trimmed, mapped, and summarized at the gene-level using the same tools as described above. A processed gene count matrix was obtained from Zhang et al. Gene count matrices were used as the input for Seurat (V3;(Butler et al., 2018; Stuart et al., 2019)) where samples were filtered, scaled, and clustered.

### Meta analysis of datasets

Individual Seurat objects were imported to R and processed using Seurat following the “Multiple Dataset Integration and Label Transfer” vignette available on the program website (https://satijalab.org/seurat/v3.1/integration.html). Cluster markers of the integrated dataset were calculated from the ‘RNA’ assay of the Seurat object using a logistic regression test.

### Cluster marker correlation analysis

Cluster markers identified in Seurat were compared for overlapping markers across analyses using a custom R script. Correlation for each comparison was calculated by dividing the number of overlapping markers by the number of published cluster markers. See Table 1 of this manuscript and Table S2 of Zhang et al 2019 for cluster markers used.

**Table 1.** Confirmation of cluster markers identified by Dvoryanchikov et al. 2017. Seurat cluster represents the cluster each marker was found in the reanalyzed data. Confirmed cluster is whether or not the cluster marker was found in the most-correlated cluster. Some markers were significant by p-value but not FDR-adjusted p-value, as denoted in the final column.

| Dvoryanchikov cluster | Gene | Seurat cluster | Confirmed marker | p <0.05, FDR-adjusted p > 0.05 |
| --- | --- | --- | --- | --- |
| Somatosensory | Fxyd2 | D0 | yes |  |
| Phox2b- | Kcns3 | D0 | yes |  |
|  | Prrxl1 | D0 | yes |  |
|  | Ldb2 | D0 | yes |  |
|  | Pdlim1 | D0 | yes |  |
|  | Mef2c | D0 | yes |  |
| TI | Olfm3 | D1,D2,D3 | yes | * (D1,D2,D3) |
| Phox2b+ | Itm2a | D2 | yes | D2 |
|  | Hspb3 | na | no |  |
|  | Foxg1 | D1,D2 | Yes | * (D1,D2) |
|  | Irf6 | D2,D3 | yes | * (D2,D3) |
|  | Eya2 | D3 | yes |  |
| TII | Cd302 | D4 | yes | * |
| Phox2b+ | Kcnip1 | D4 | Yes | * |
|  | Slc38a11 | D4 | yes |  |
|  | Mafb | D4 | yes | * |
|  | Meis1 | D4 | Yes | * |
|  | Tbx2 | - | no |  |
|  | Dach2 | D4 | Yes |  |
|  | Runx3 | - | No |  |
| TIII | Lypd1 | D5 | yes |  |
| Phox2b+ | Penk | D5 | yes |  |
|  | Trhr | D5 | yes |  |
|  | Prox2 | - | no |  |
|  | Phox2a | - | no |  |
|  | Twist2 | - | no |  |

### Data visualization

Data were visualized using the pheatmap package in R or with Seurat plotting features DimPlot and DoHeatmap and FeaturePlot (Butler et al., 2018; Kolde, 2019; Stuart et al., 2019).

## Results

### Characterization of Phox2b-cre mouse

To confirm that the *Phox2b*-cre driver faithfully drives CRE recombinase in PHOX2B-expressing cells, geniculate ganglia and gustatory taste fields of *Phox2b*-cre/Rosa-tomato mice were examined using confocal immunofluorescence microscopy. A subset of geniculate ganglion neurons expresses robust tdTomato fluorescence consistent with previous reports (Ohman-Gault et al., 2017; Rios-Pilier and Krimm, 2019). The majority of tdTomato-expressing neurons exhibited P2X3 immunoreactivity, and conversely, nearly all P2X3-immunoreactive neurons exhibited tdTomato fluorescence (Figure 1). To confirm that tdTomato is being driven in PHOX2B-expressing ganglion cells, sections were labeled with an antibody against PHOX2B. Indeed, every tdTomato-expressing neuron showed positive nuclear labeling of the transcription factor (Figure 1).

**Figure 1.**
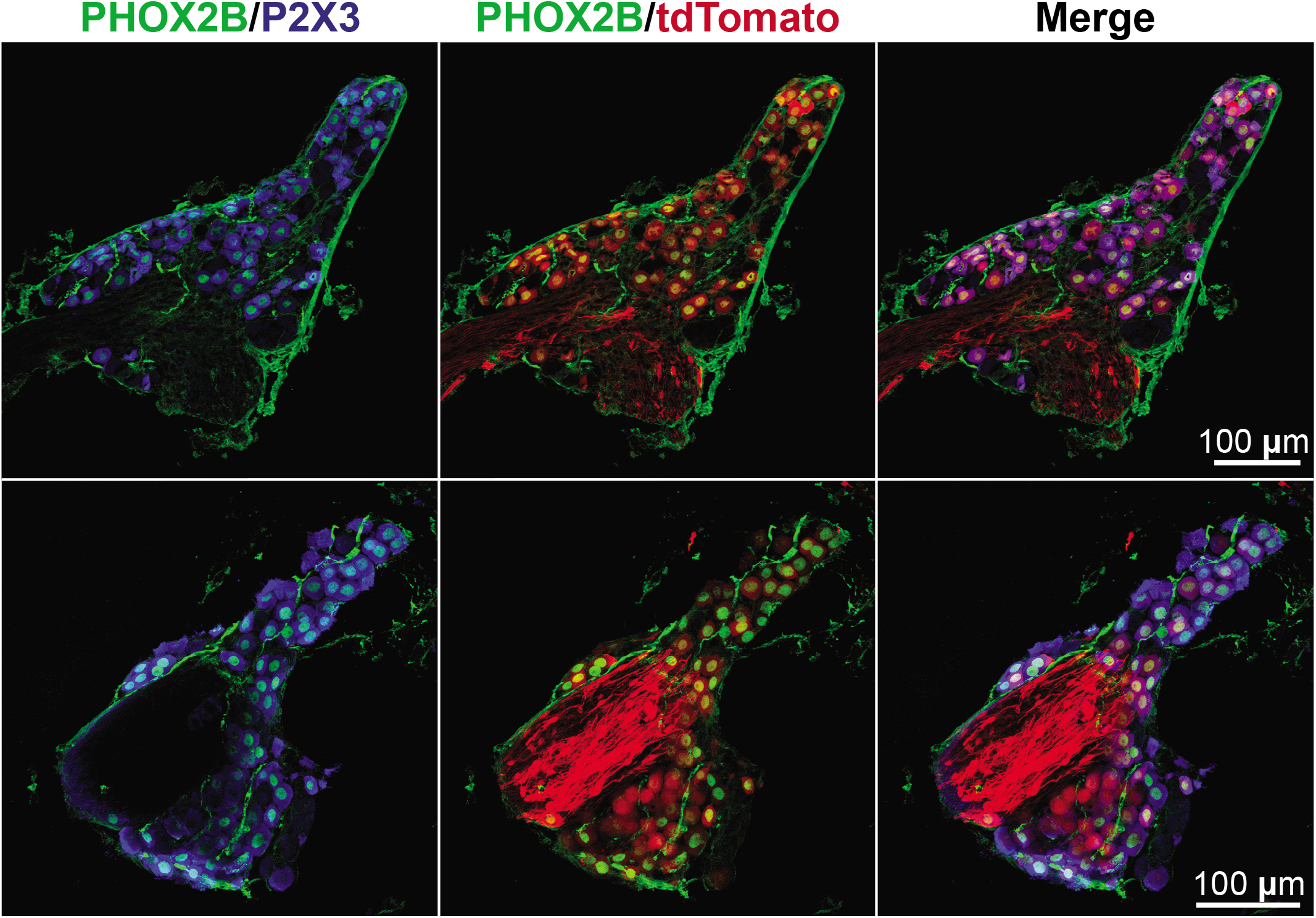
tdTomato expression in PHOX2B-immunoreactive neurons. Sections of Phox2bcre/tdTomato geniculate ganglion labeled with antibodies against PHOX2B and P2X3. Nearly all tdTomato-expressing cells are immunoreactive for PHOX2B and P2X3. Non-nuclear “green” signal is due to binding of anti-mouse secondary antibodies to endogenous mouse IgG; this signal persists in no primary controls, and no nuclear signal is observed. Sections are maximal z-projections of ~12 μm image stacks.

Within taste papillae, tdTomato positive nerve fibers enter taste buds consistent with the *Phox2b* as a marker of gustatory nerve fibers. Most tdTomato-expressing nerve fibers exhibit strong P2X3-immunoreactivity. Additionally, these fibers are rarely outside of taste buds but when present are not immunoreactive for P2X3 (Figure 2)

**Figure 2.**
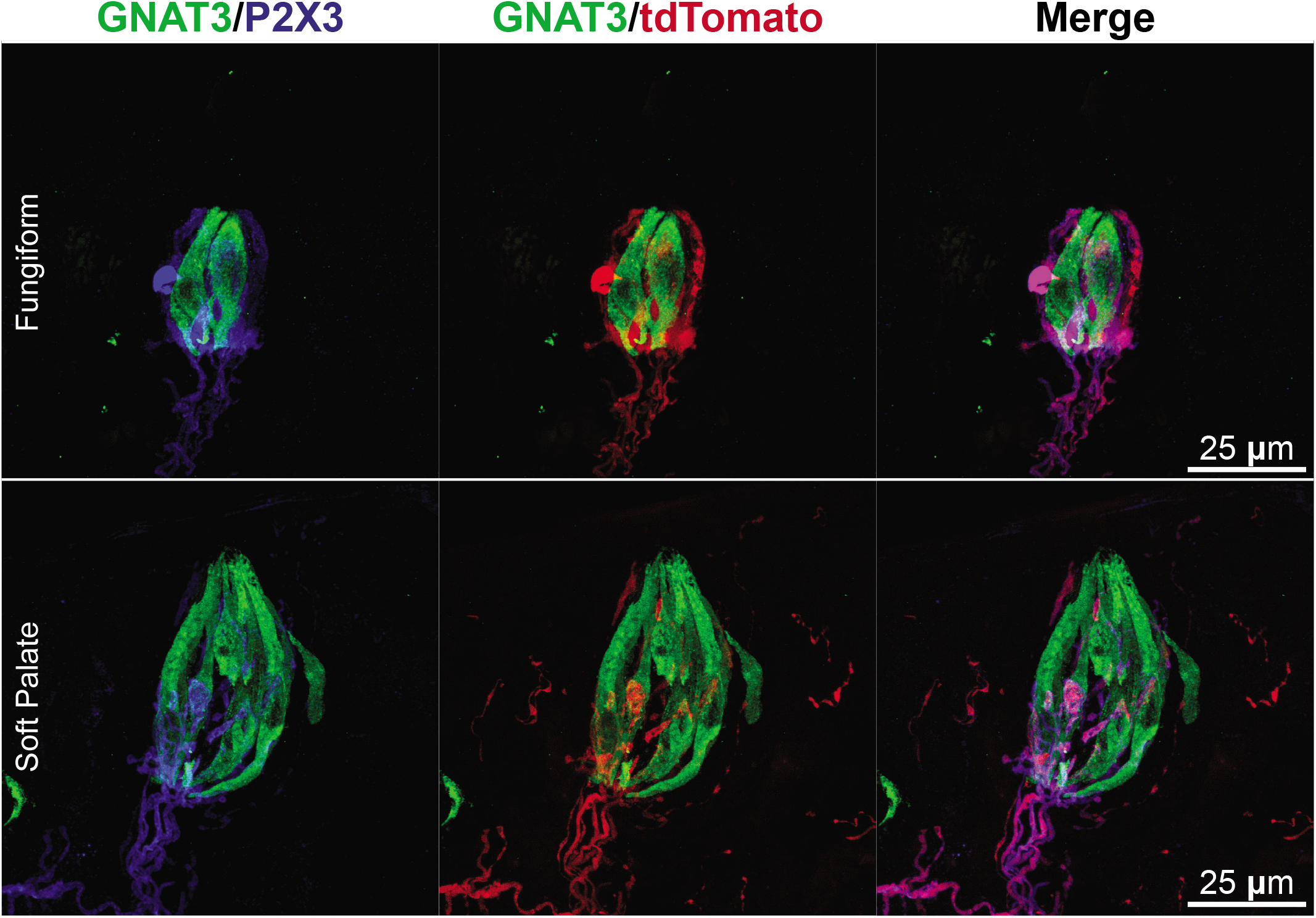
tdTomato-expressing nerve fibers enter taste buds. Sections of lingual taste papillae showing immunoreactivity to GNAT3 (α-gustducin; Type II cells) and P2X3 with endogenous tdTomato fluorescence. Within taste buds, all tdTomato+ fibers exhibit P2X3 immunoreactivity. In some cases, tdTomato+ fibers were found in the intergemmal area and were P2X3 negative. Images are maximal z-projections of ~12-16 μm image stacks.

### scRNA-seq of Phox2b geniculate ganglion neurons

Single cell RNA sequencing was performed on manually isolated geniculate ganglion neurons (48 samples from 6 total mice across 2 collection days; 40 tdTomato+, 6 tdTomato-, 2 blanks). Ganglion neurons were enzymatically digested and plated on a Qiagen QiaScout Microraft array. Using the QiaScout device cells were visually inspected for presence or absence of tdTomato, dislodged from the microraft array, and transferred to cell lysis buffer where lysis was initiated followed by reverse transcription, whole transcriptome amplification, and sequencing library preparation. Post processing of the sequencing data excluded samples that had < 50% mapping rate, expression of *Snap25* or *Uchl1* < 5 TPM, and <2000 unique genes. Post processing resulted in 31 cells (17 tdTomato+, 4 tdTomato-). *Phox2b* expression correlated with tdTomato fluorescence (Figure 3) and taste-related transcripts (*P2rx2*, *P2rx3*) were present in all *Phox2b*+ neurons. A subset of *Phox2b*+ neurons also expressed *Htr3a* (Figure 3). Additionally, genes such as *Penk and Spon1*, markers of sour-best and sweet-best neurons (Zhang 2019) were identified in some neurons. Unfortunately, the low sample size prevented use of unbiased clustering algorithms such as Seurat (Stuart et al., 2019).

**Figure 3.**
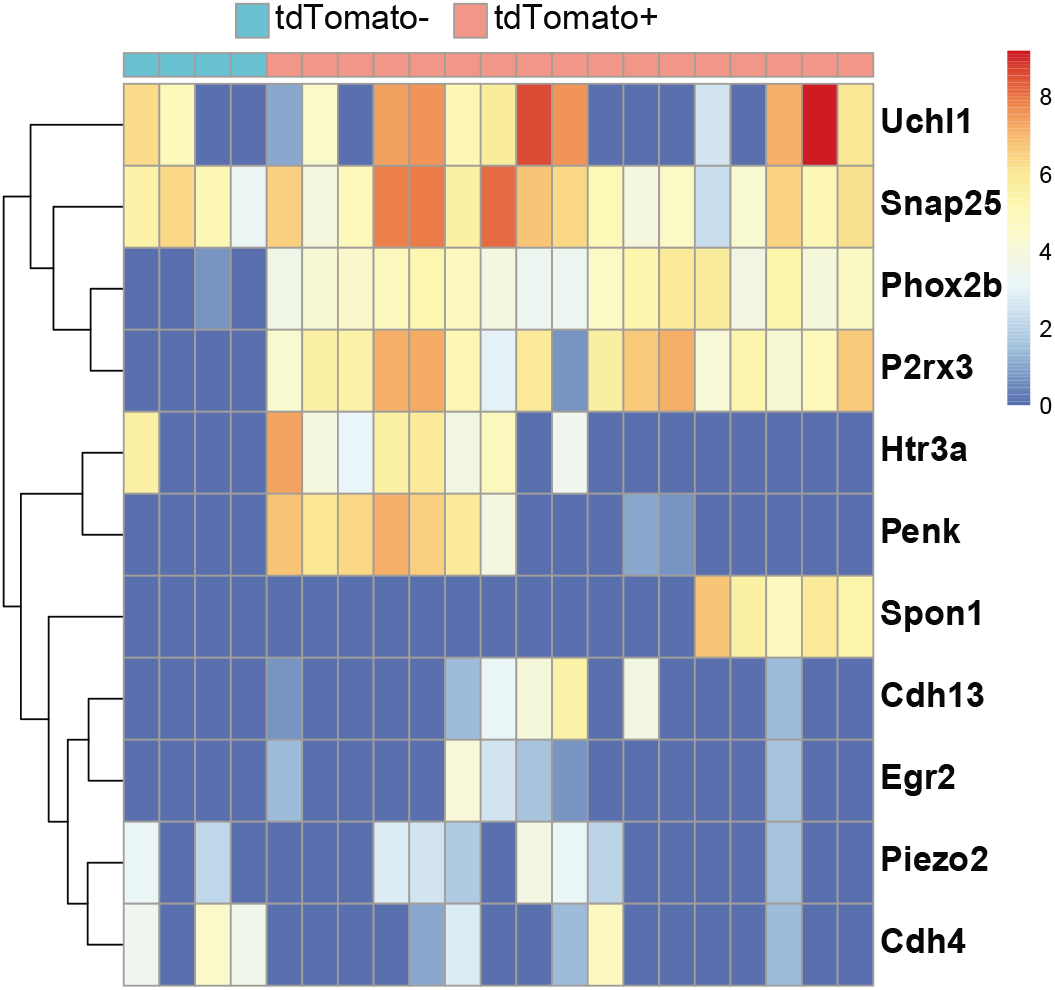
Expression of select transcripts in tdTomato-expressing neurons. Geniculate ganglion neurons were collected using the QiaScout and processed for single cell RNA sequencing using the Qiagen FX single cell RNA kit. Heatmap shows log(transcripts per million) of select transcripts. *Phox2b* expression correlates with presence or absence of tdTomato.

### Replication of published results

The first report of scRNA-seq on geniculate ganglion neurons used wildtype mice and the Fluidigm C1 platform to perform expression profiling and demonstrated 4 cell clusters (3 gustatory, 1 non-gustatory). Reanalysis of these 96 cells using Seurat reveals 6 clusters of cells (Figure 4A,B) including 1 somatosensory (Phox2b-) and 5 gustatory (Phox2b+) clusters. Cluster markers described by Dvoryanchikov et al. readily mark clusters in the reanalyzed data (Figure 4C). Correlation analysis of cluster markers from the original study and the reanalyzed data show strong concordance of data (Figure 4D). However, the main difference is that in the reanalyzed data, three clusters (D1,D2,D3) correlate with the original TI cluster. We attribute the discrepancy among cluster numbers likely arises due to 1) different clustering software/algorithms and 2) less conservative clustering variables. Overall, we show that this independent analysis of data from Dvoryanchikov et al. yields similar cell clustering results. See Tables 1 and 2 for specific cluster markers.

**Figure 4.**
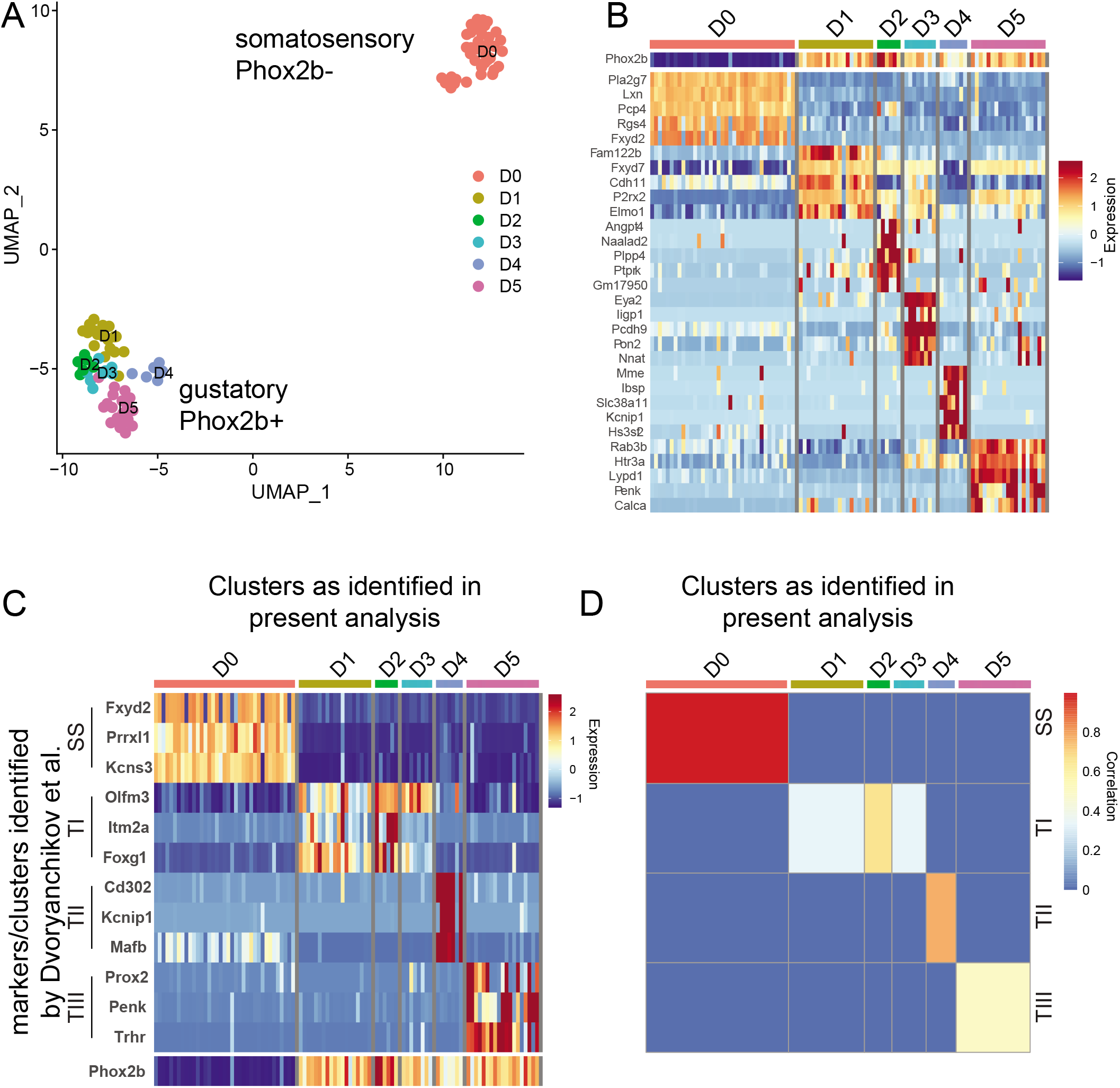
Secondary analysis of Dvoryanchikov et al. 2017 scRNA-seq data. Analysis of cells publicly available from GSE102443. A. Seurat UMAP clustering of cells expressing greater than 2000 genes. B. Heatmap showing transcript expression of top 5 cluster markers identified using Seurat. C. Heatmap showing transcript expression of cluster markers described in Dvoryanchikov et al. 2017. Note, that the Dvoryanchikov et al 2017 cluster T1 is further subdivided into 3 sub-clusters identified here as D1, D2, and D3. D. Correlation of cluster markers described by Dvoryanchikov et al. 2017 and those identified in this study using Seurat.

**Table 2.** Top cluster markers from reanalysis of Dvoryanchikov et al 2017. Markers ordered by ascending FDR-adjusted p value

| cluster |  |  |  |  |  |
| --- | --- | --- | --- | --- | --- |
| Phox2b- | Phox2b+ |  |  |  |  |
| D0 | D1 | D2 | D3 | D4 | D5 |
| Tafa3 | Fam122b | Angpt4 | Eya2 | Mme | P2ry12 |
| Hs6st2 | Fxyd7 | Naalad2 | Ligp1 | Dach2 | Creg2 |
| Antxr2 | Pxmp2 | Plpp4 | Pcdh9 | Ibsp | Pdlim3 |
| Mob3b | Unc5c | Ptprk | Pon2 | Draxin | Fstl4 |
| Prmt8 | Rassf9 | Gm17950 | Nnat | Col11a1 | Rab3b |
| Hoxd1 | Bambi | Kcnk2 | Plxdc2 | Slc38a11 | Trhr |
| Nrp1 | Thsd7b | Trappc3l | Gnas | Kcnip1 | Htr3a |
| Ldb2 | Cdh11 | Spon1 | Caprin1 | Meis1 | Sorbs2 |
| Prrxl1 | P2rx2 | Flrt2 | Cav1 | Kif21b | Rgs17 |
| Syt6 | Foxg1 | Fkbp11 | Cd34 | Hs3st2 | Trim25 |

We next applied a similar processing pipeline to scRNA-seq data from Zhang et al 2019. The authors manually collected over 800 *P2rx3*-tdTomato expressing geniculate ganglion cells and performed scRNA-sequencing using the Cel-seq technology. Of the 454 neurons available from GEO: GSE135801, 372 had expression of >2000 unique genes. Zhang et al originally reported 7 total clusters (labeled A-G). Gustatory clusters (*Phox2b*+) were identified by *Spon1*, *Penk*, *Cdh4*, *Cdh13*, or *Egr2*. Reanalysis of these data using Seurat reveals 7 molecularly distinct neuronal clusters (Z0-Z6) including those marked by *Spon1* (cluster Z2), *Penk* (cluster Z6), *Cdh13* (cluster Z1), *Egr2* (cluster Z3) and *Piezo2* (clusters Z0 and Z5; Figure 5B,C). However, *Cdh4* was not detected as a significant marker and *Cdh13* was not significant by FDR-adjusted p value (see Table 3 and 4 for a more comprehensive list of cluster markers). While these specific cluster markers were less consistent between the original report and the reanalyzed data, pairwise correlation between original cluster markers (A-G) and reanalyzed cluster markers (Z0-Z6) show that the reanalyzed clusters show overall agreement with the original report (Figure 5D). Clusters Z1 and Z4 do not strongly correlate with any of the clusters from the original report. Likewise, clusters A and B do not correlate strongly with any clusters identified in the reanalyzed data. Interestingly, cells belonging to cluster Z4 had the least unique genes detected per cell, the least number of reads per cell, and the fewest significant cluster markers (not shown). Thus cluster Z4 could represent a cluster of unhealthy cells and/or poorly sequenced cells. Interestingly, when the data are reprocessed after removing cells with less than 4000 unique genes detected (as opposed to 2000), cluster Z4 disappears with overall preservation of the other clusters (not shown).

**Figure 5.**
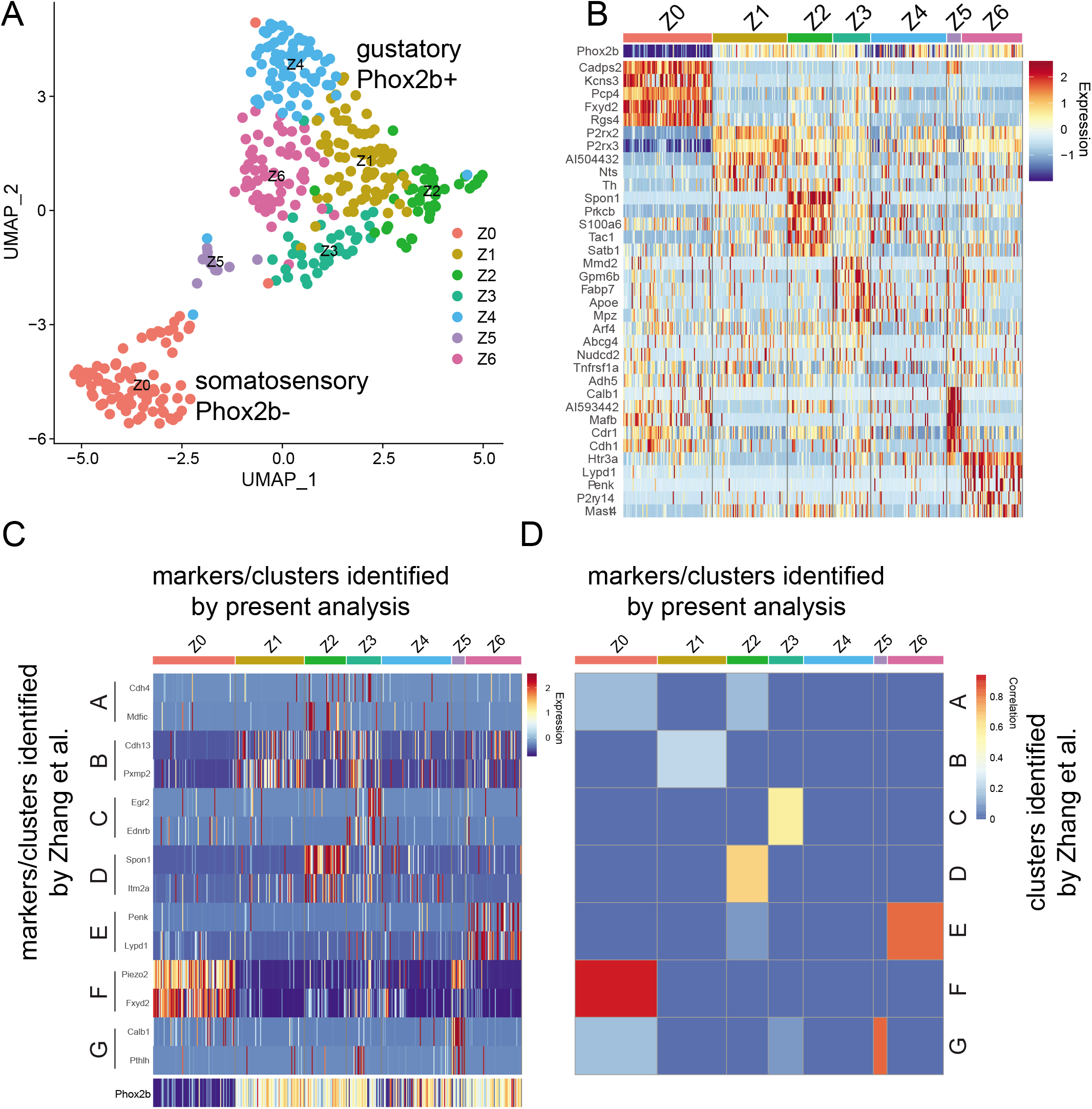
Secondary analysis of Zhang et al. 2019 scRNA-seq data. Analysis of cells publicly available from GSE135801. A. Seurat UMAP clustering of cells expressing greater than 2000 genes. B. Heatmap showing transcript expression of top 5 cluster markers identified using Seurat. C. Heatmap showing transcript expression of cluster markers described in Zhang et al. 2019. D. Correlation of cluster markers described by Zhang et al. 20179 and those identified in this study using Seurat.

**Table 3.**
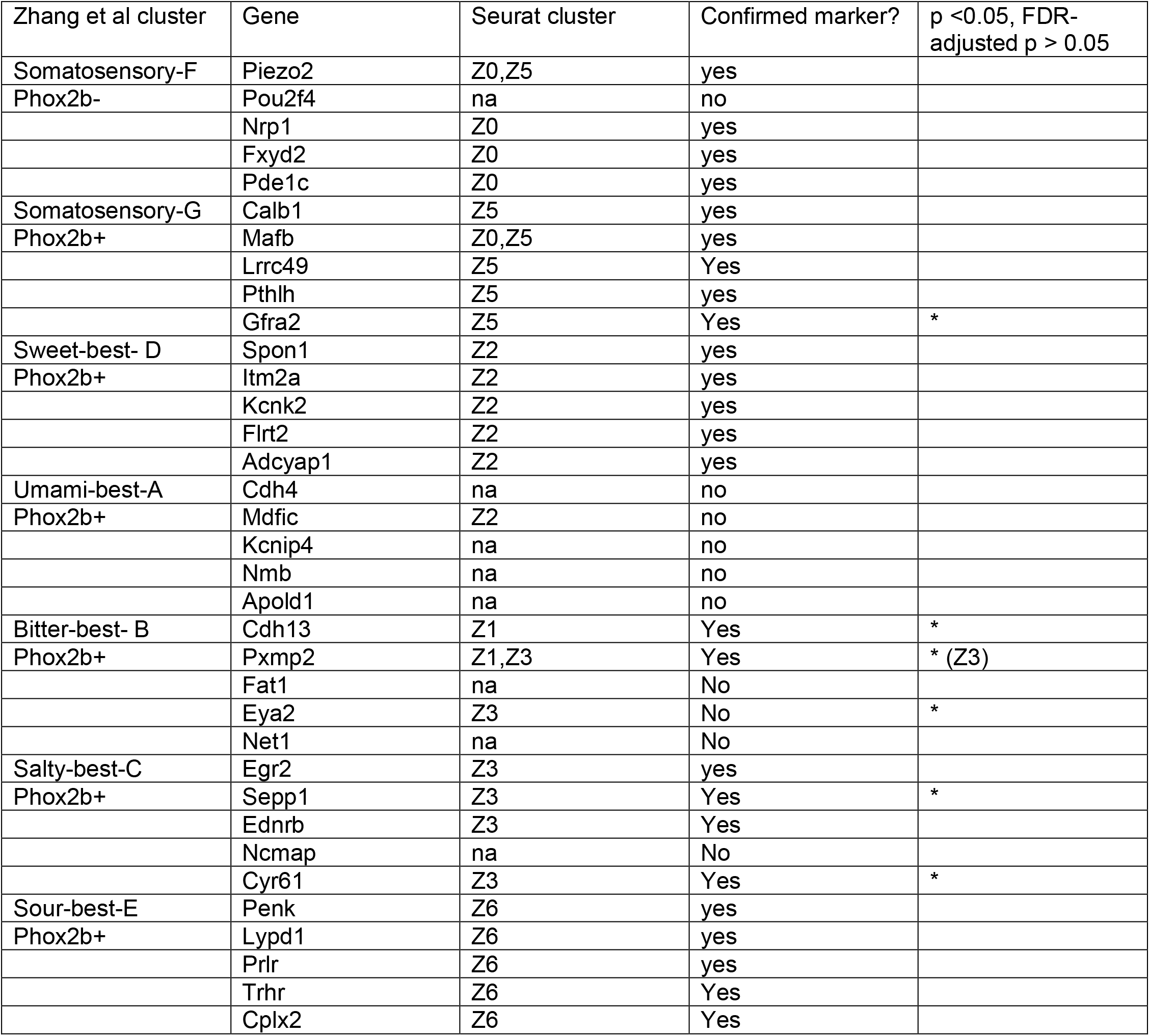
Confirmation of cluster markers identified by Zhang et al. 2019. Seurat cluster represents the cluster each marker was found in the reanalyzed data. Confirmed cluster is whether or not the cluster marker was found in the most-correlated cluster. Some markers were significant by p-value but not FDR-adjusted p-value, as denoted in the final column.

**Table 4.**
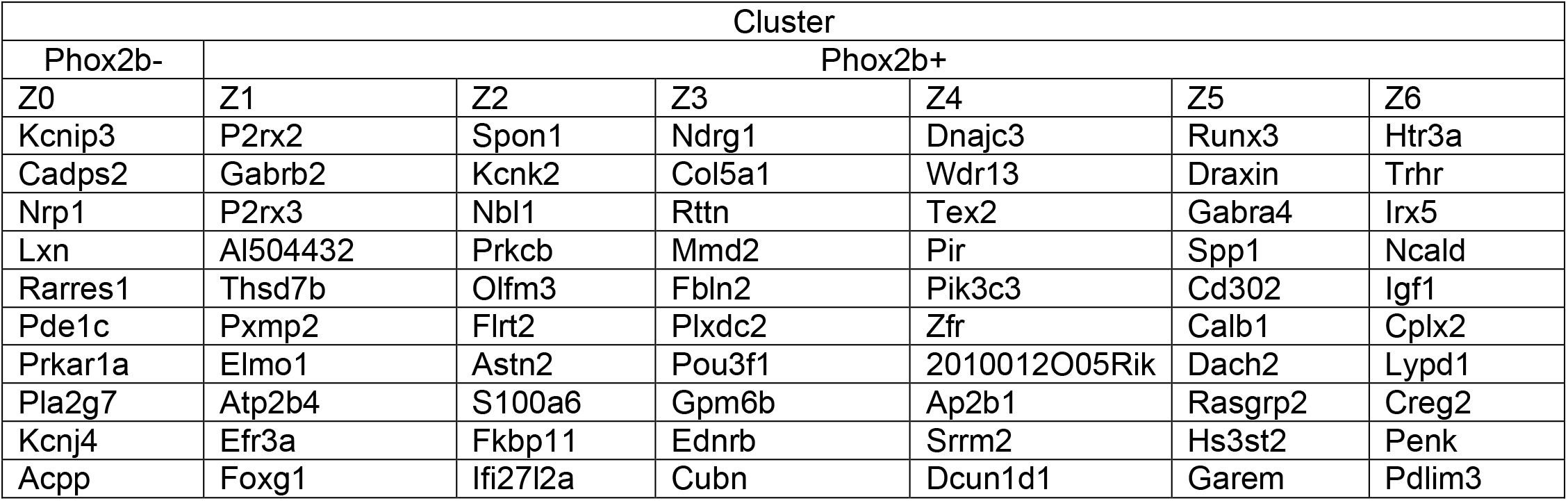
Top cluster markers from reanalysis of Zhang et al 2018. Markers ordered by ascending FDR-adjusted p value

**Table 5.** Comparison across datasets.

| Cluster from combined analysis | corresponding cluster from Zhang et al | Corresponding cluster from Dvoryanchikov et al. | Corresponding cluster from Zhang This analysis | Corresponding cluster from Dvoryanchikov This analysis | Potential Function |
| --- | --- | --- | --- | --- | --- |
| M0 | F- Somatosensory (Piezo2) | Somatosensory | Z0 | D0 | Somatosensory |
| M1 | B- Cdh13/pxmp2 cluster | TI | Z1 | D1 |  |
| M2 | D- Spon1 cluster | TI | Z2 | D2 | Sweet-best |
| *M3* | *B- Cdh13/pxmp2 cluster* | *TI* | *Z3* | *D3* |  |
| *M4* |  |  | *Z4* |  | *Poor quality cells?* |
| M5 | G- Somatosensory2 (Piezo2) | TII | Z5 | D4 | Somatosensory? |
| M6 | E- Penk cluster | TIII | Z6 | D5 | Sour-best |

### Comparison across datasets

Overall, independent reanalysis of both datasets using a parallel analysis pipeline agrees with the original reports. We next asked whether clusters from one dataset correlate to the other dataset. First, we examined how top cluster markers identified by the original reports correlate with cluster markers identified during reanalysis of the other dataset (ie do the cluster markers identified by Zhang et al 2019 correlate with cluster markers identified by reanalysis of Dvoryanchikov et al 2017?). Cluster markers identified by Zhang et al 2019 correlate strongly with some, but not all, markers identified by reanalysis of Dvoryanchikov et al 2017 (Figure 6A). For instance, clusters D0, D2, D4, and D5 from reanalysis of Dvoryanchikov et al 2017 correlate strongly with clusters F, D, G, and E of Zhang et al 2019, respectively. Conversely, cluster markers identified by Dvoryanchikov et al 2017 also correlate strongly with most markers identified by reanalysis of Zhang et al 2019 (Figure 6B). Clusters Z0, Z1, Z2, Z5, and Z6 from reanalysis of Zhang et al 2019 correlate strongly with clusters somatosensory, TI, TI, TII, and TIII of Dvoryanchikov et al 2017, respectively. From these comparisons, we can conclude with higher confidence that certain clusters are present and valid as they are seen across both datasets and contain a high proportion of overlapping markers identified by all analysis. The clusters identified in the original reports that we have the most confidence in (in descending order) are the somatosensory cluster (SS, F, D0, Z0), *Penk*-expressing gustatory cluster (TIII, E, D5, Z6), *Calb1*-expressing gustatory cluster (TII, G, D4, Z5), *Spon1*-expressing gustatory cluster (TI, D, D2, Z2), and Pxmp2-expressing gustatory cluster (TI, B, Z1, D1). The remaining clusters show less correlation to the original reports. Next, we compared the cluster markers identified by independent reanalysis of both datasets to find overlapping markers. Indeed, there was a degree of concordance amongst the clusters (Figure 6C). We conclude that 5 clusters from each dataset share many overlapping markers and thus are likely the same identity.

**Figure 6.**
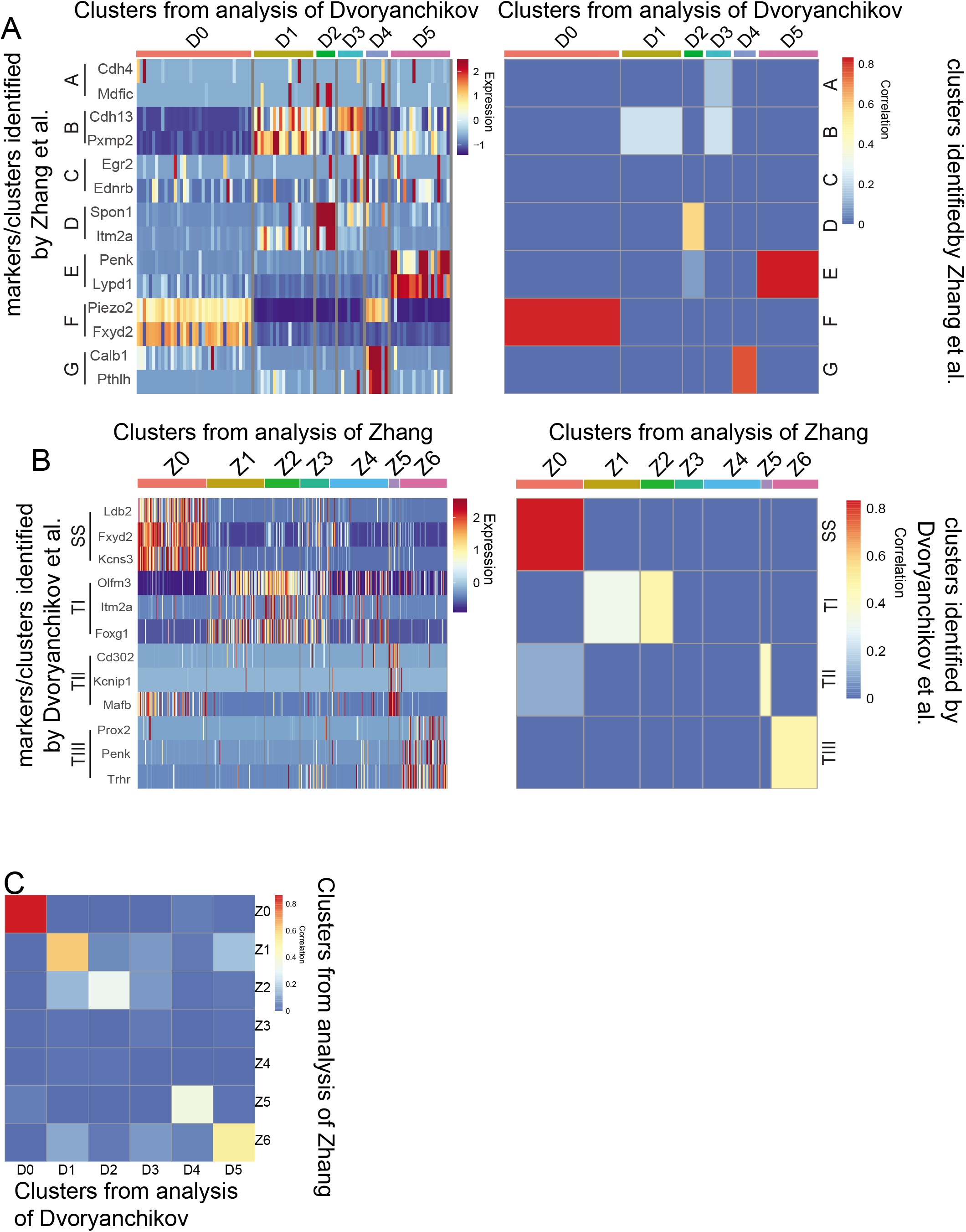
Comparison of published cluster markers and cluster markers identified from reanalysis of both datasets. A. Heatmap showing reanalyzed data from Dvoryanchikov et al 2017 with markers identified in the original report of Zhang et al 2019. Correlation heatmap shows pairwise comparisons between cluster markers. B. Heatmap showing reanalyzed data from Zhang et al 2019 with markers identified in the original report of Dvoryanchikov et al 2017. Correlation heatmapshows pairwise comparisons between cluster markers. C. Correlation heatmap of cluster markers from both reanalyzed datasets

### Meta analysis of all datasets

Generally, in RNA-sequence analysis, it is against best-practices to combine datasets from multiple sources given the range of variability in sample preparation, sequencing, and data processing. However, with the recent release of Seurat V3 a method for combining datasets from single cell sequencing experiments across multiple technologies has been developed (Stuart et al., 2019). This method finds integration anchors common amongst the datasets, merges the data, and scales the data based on the anchors. This meta analysis allows for integration of data on similar cell types from multiple sources.

The three datasets in this paper can be combined, scaled, and clustered together. Interestingly, UMAP plotting of the integrated data shows a strong similarity among the three datasets, with cells from each dataset being represented throughout the plot (Figure 7A,B). This suggests that despite three different methods and technologies used to collect scRNA-seq data, the genetic information was similar between studies. Clustering of the integrated data reveals 7 cell clusters. These clusters largely share markers identified by both the original reports and the individually reanalyzed datasets. With the exception of cluster M4, all identified clusters from the combined dataset show overlapping markers with clusters from the separate datasets (Figure 7F,G, Table 6). For reasons discussed above, cluster M4 likely represents a population of unhealth of poorly sequenced cells. When the datasets are reintegrated after removing cells with less than 4000 unique genes detected, cluster M4 disappears with overall preservation of the other clusters (not shown).

**Figure 7.**
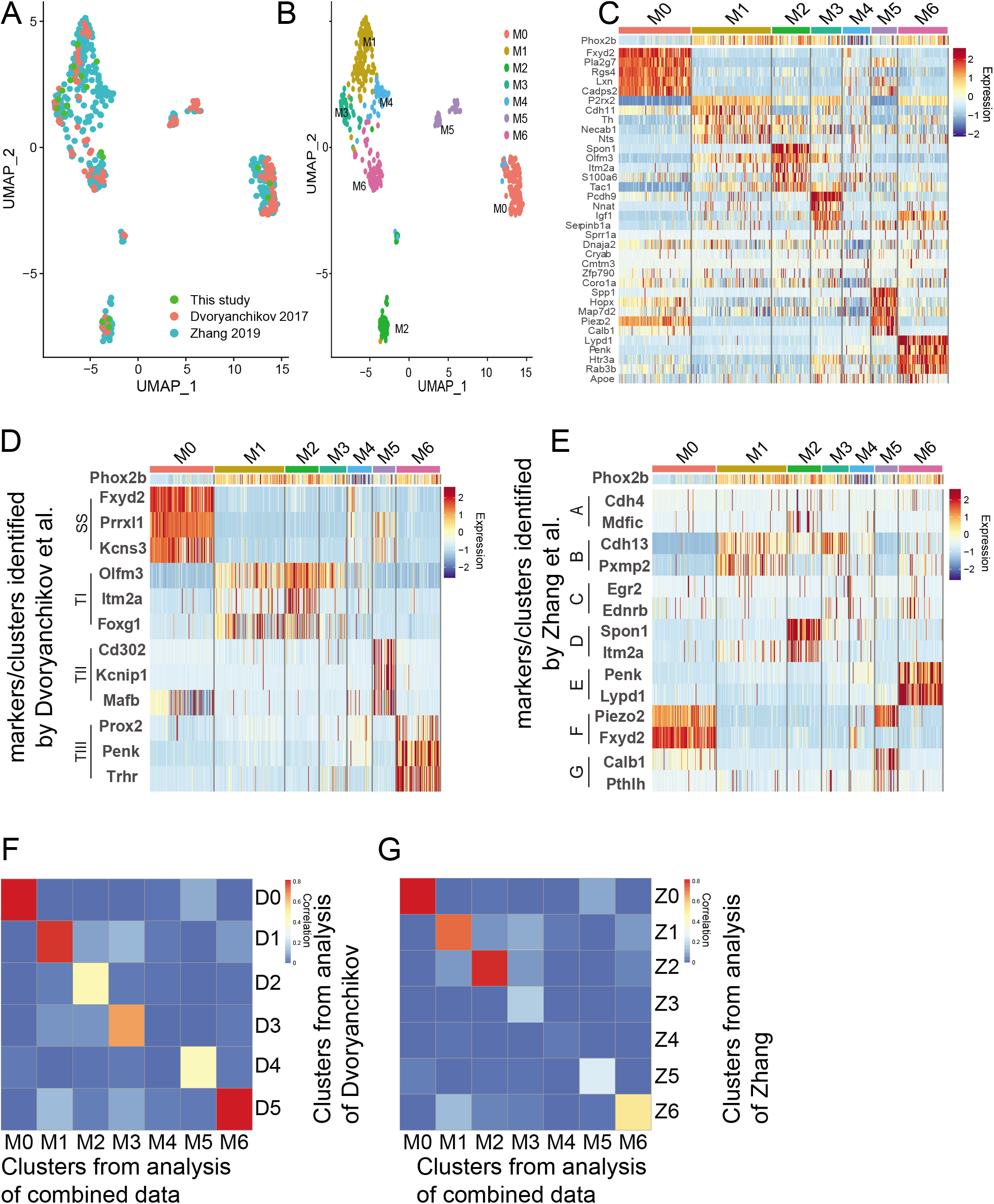
Combination of data from this paper, Dvoryanchikov et al 2017, and Zhang et al 2019. A. Seurat UMAP clustering showing each dataset. B. Overall clusters of the combined datasets. C. Heatmap showing transcript expression of top 5 markers of each cluster. D,E. Heatmaps of combined dataset analysis showing expression of cluster markers identified in the original report of Dvoryanchikov et al 2017 (D) and Zhang et al 2019 (E). F,G. Correlation heatmap of cluster markers from the combined dataset analysis versus cluster markers identified by reanalysis of Dvoryanchikov et al 2017 (F) or Zhang et al 2019 (G).

**Table 6.**
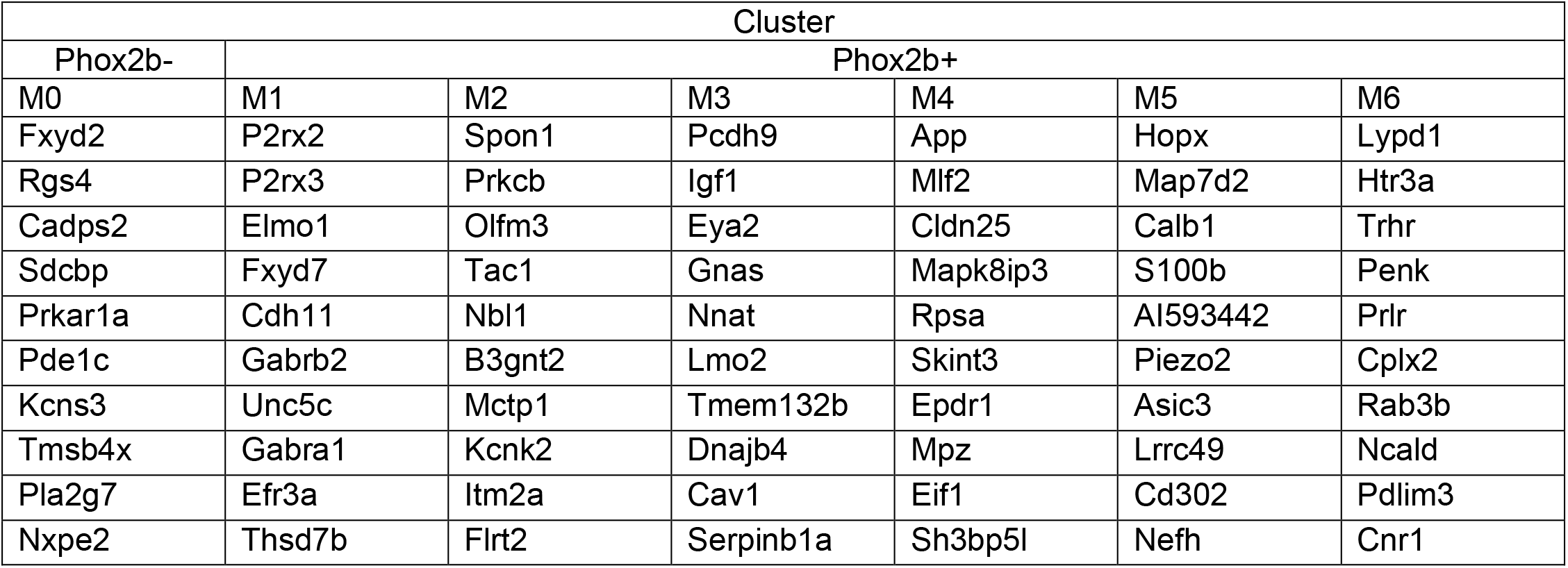
Top Cluster Markers combined dataset.

## Discussion

### Do neuron clusters represent specific taste modalities?

Single cell RNA-seq allows for the determination of heterogeneity within a cell population. By clustering cells based on similarities, markers that are unique to those clusters can be identified, which can give strong insight to their function. While Dvoryanchikov et al. were the first to reveal molecular heterogeneity by identifying clusters of geniculate ganglion neurons, Zhang et al. were the first to attribute the role of specific cell clusters to taste detection. Convincing evidence demonstrates that neurons in the *Penk* and *Spon1*-expressing cell clusters represent sour- and sweet-best neurons. Thus, it is plausible that these neurons receive specific input from Type III and Type II taste receptor cells, respectively.

Sour and sweet: All three datasets analyzed in the present study showed expression of *Penk* and *Spon1* in gustatory fibers and that they marked specific cell clusters. In Zhang et al 2019, *Penk* and *Spon1* were cluster markers. Dvoryanchikov et al 2017 reported *Penk* as a cluster marker, but not *Spon1*. Interestingly, secondary analysis reveals that *Spon1* neurons were a subset of the cluster labeled ‘T1’ in the original report (Dvoryanchikov et al. 2017), which was subdivided in to three clusters in this current study (Figure 5C). These findings together with the in vivo imaging of Zhang et al 2019 strongly suggest that sour-best and sweet-best fibers are molecularly distinct from other ganglion neurons.

Bitter, salty, and umami: Zhang et al reports that *Cdh4*, *Egr2*, and *Cdh13* are cluster markers for bitter, salty, and umami-best neurons. However, in the present analyses, these markers did not define clusters as well as *Penk* and *Spon1*. Additionally, the behavioral assays performed by Zhang et al relied on constitutive knockout of *Cdh4* and *Cdh13* and tetanus toxin inactivation of *Egr2* neurons. While the experiments showed that umami, bitter, and salty taste preference rely on these transcripts, they do not show that this requirement is at the level of the geniculate ganglion. Indeed, according to the Allen Brain Map, these three transcripts are expressed in a variety of cell types throughout the central nervous system. Additionally, according to secondary analysis of Qin et al, *Cdh13* is expressed in both Type II and Type III taste receptor cells and *Egr2* is expressed in some Type III cells (not shown; Qin et al., 2018). Thus, any ganglion cell clusters responsible for the detection of bitter, salty, and umami signals from taste receptor cells remains enigmatic.

### Does Htr3a mark a specific cluster?

Stratford et al 2017 previously showed that nerve fibers expressing the serotonin receptor 5-HT3 preferentially contact Type III taste receptor cells in all taste fields (Stratford et al 2017) suggesting that these fibers may be molecularly distinct within the geniculate ganglion. However, Dvoryanchikov et al. argued that *Htr3a* did not delineate a specific cluster in their analyses. In the present analyses, all three datasets show clustering of Htr3a-expressing neurons. While *Htr3a* was in multiple clusters, it had strongest expression in the cluster marked by *Penk* – the sour-best fibers which support our findings that these cells wire to sour-sensing Type III taste receptor cells. Conversely, *Htr3a* is absent in the *Spon1* cluster (sweet-best fibers). Interestingly, in all datasets, *Htr3a* is also present in a gustatory (*Phox2b*+) cluster expressing *Piezo2*, a transcript for a mechanosensitive channel. This cluster expressed low levels of *P2rx3*, and no *P2rx2*. Moayedi et al 2018 showed that *Piezo2*-GFP expressing nerve fibers are present in fungiform papillae, just adjacent to taste buds, and that these fibers enter the papillae in a bundle with other gustatory fibers (Moayedi et al., 2018). Interestingly, P2X2/3 double knockout mice (which lack nerve response to all tastants) still show nerve responses to mechanical stimulation of the tongue (Finger et al., 2005). Perhaps this *Piezo2*/*Htr3a*/*Phox2b* neuronal cluster is responsible for this ATP-independent response.

### Uncorrelated clusters

We report that across datasets, there are some clusters that do not correlate with any other clusters (primarily Z3, and Z4, and D3; Figure 6C). One possibility, as discussed above, is that some of these clusters could represent unhealthy or poorly sequenced cells. This is primarily the case with cluster Z4. For the remaining clusters, it could be that the sample size of the studies may not be large enough to encompass all neuron clusters. It could be that each uncorrelated cluster represents a different cluster of cells. Thus, larger geniculate ganglion scRNA-seq databases must be obtained.

## Conclusions

Overall, the data analyzed in this paper confirm molecular heterogeneity among the neurons of the geniculate ganglion. The datasets clearly separate gustatory vs non-gustatory neurons by expression of *Phox2b*. Within the *Phox2b* expressing fibers, clear sub clusters are formed. Consistent among the datasets are clusters defined by *Penk*, *Spon1*, and *Piezo2*. Zhang et al showed that *Penk* clusters represent sour-best fibers while *Spon1* clusters represent sweet-best fibers. The *Piezo2* cluster could be responsible for residual chorda tympani nerve responses to lingual mechanical stimuli in the absence of purinergic signaling (Finger et al. 2005). The remaining clusters were more variable between the datasets and markers identified by Zhang et al (*Cdh4*, *Cdh13*, *Egr2*) were not clearly observed in this study. While it is likely that neurons responsible for detecting each taste modality are molecularly distinct, more work needs to be done to elucidate the molecular phenotypes of bitter, salty, and umami sensitive fibers.

## Figure Legends

Table S1. Seurat-identified cluster markers for Zhang et al 2018.

Table S2. Seurat-identified cluster markers for Dvoryanchikov et al 2017.

Table S3. Seurat-identified cluster markers for meta analysis.

## Acknowledgments

We thank Qiagen for allowing us to demo the QiaScout device and supplying samples of the QiaScout Microraft Arrays. We also thank Drs. Tom Finger and Sue Kinnamon for their support and critique of the manuscript.

## References

Bushnell, B. BBMap.

Butler, A., Hoffman, P., Smibert, P., Papalexi, E., and Satija, R. 2018. Integrating single-cell transcriptomic data across different conditions, technologies, and species. Nature Biotechnology. 36:411–420.

Dvoryanchikov, G., Hernandez, D., Roebber, J.K., Hill, D.L., Roper, S.D., and Chaudhari, N. 2017. Transcriptomes and neurotransmitter profiles of classes of gustatory and somatosensory neurons in the geniculate ganglion. Nature Communications. 8:760.

Finger, T.E., Danilova, V., Barrows, J., Bartel, D.L., Vigers, A.J., Stone, L., Hellekant, G., and Kinnamon, S.C. 2005. ATP Signaling Is Crucial for Communication from Taste Buds to Gustatory Nerves. Science. 310:1495–1499.

Hashimshony, T., Wagner, F., Sher, N., and Yanai, I. 2012. CEL-Seq: Single-Cell RNA-Seq by Multiplexed Linear Amplification. Cell Reports. 2:666–673.

Huang, Y.-J., Maruyama, Y., Dvoryanchikov, G., Pereira, E., Chaudhari, N., and Roper, S.D. 2007. The role of pannexin 1 hemichannels in ATP release and cell–cell communication in mouse taste buds. PNAS. 104:6436–6441.

King, M.S., and Bradley, R.M. 2000. Biophysical properties and responses to glutamate receptor agonists of identified subpopulations of rat geniculate ganglion neurons. Brain Research. 866:237–246.

Koga, T., and Bradley, R.M. 2000. Biophysical Properties and Responses to Neurotransmitters of Petrosal and Geniculate Ganglion Neurons Innervating the Tongue. J Neurophysiol. 84:1404–1413.

Kolde, R. 2019. pheatmap: Pretty Heatmaps.

Larson, E.D., Vandenbeuch, A., Voigt, A., Meyerhof, W., Kinnamon, S.C., and Finger, T.E. 2015. The Role of 5-HT3 Receptors in Signaling from Taste Buds to Nerves. J Neurosci. 35:15984–15995.

Malin, S.A., Davis, B.M., and Molliver, D.C. 2007. Production of dissociated sensory neuron cultures and considerations for their use in studying neuronal function and plasticity. Nat Protocols. 2:152–160.

Moayedi, Y., Duenas-Bianchi, L.F., and Lumpkin, E.A. 2018. Somatosensory innervation of the oral mucosa of adult and aging mice. Sci Rep. 8:1–14.

Mtui, E., Gruener, G., and FitzGerald, M. 2011. Clinical Neuroanatomy and Neuroscience - 6th Edition.

Ohman-Gault, L., Huang, T., and Krimm, R. 2017. The transcription factor Phox2b distinguishes between oral and non-oral sensory neurons in the geniculate ganglion. J Comp Neurol. 525:3935–3950.

Patro, R., Duggal, G., Love, M.I., Irizarry, R.A., and Kingsford, C. 2017. Salmon: fast and bias-aware quantification of transcript expression using dual-phase inference. Nat Methods. 14:417–419.

Qin, Y., Sukumaran, S.K., Jyotaki, M., Redding, K., Jiang, P., and Margolskee, R.F. 2018. Gli3 is a negative regulator of Tas1r3-expressing taste cells. PLOS Genetics. 14:e1007058.

R Core Team. 2019. R: A Language and Environment for Statistical Computing. Vienna, Austria: R Foundation for Statistical Computing.

Rios-Pilier, J., and Krimm, R.F. 2019. TrkB expression and dependence divides gustatory neurons into three subpopulations. Neural Dev. 14:3.

Romanov, R.A., Lasher, R.S., High, B., Savidge, L.E., Lawson, A., Rogachevskaja, O.A., Zhao, H., Rogachevsky, V.V., Bystrova, M.F., Churbanov, G.D., et al. 2018. Chemical synapses without synaptic vesicles: Purinergic neurotransmission through a CALHM1 channel-mitochondrial signaling complex. Sci Signal. 11:eaao1815.

Romanov, R.A., Rogachevskaja, O.A., Bystrova, M.F., Jiang, P., Margolskee, R.F., and Kolesnikov, S.S. 2007. Afferent neurotransmission mediated by hemichannels in mammalian taste cells. The EMBO Journal. 26:657–667.

Soneson, C., Love, M.I., and Robinson, M.D. 2016. Differential analyses for RNA-seq: transcript-level estimates improve gene-level inferences. F1000Res. 4.

Stratford, J., Larson, E., Yang, R., Salcedo, E., and Finger, T. 2017. 5-HT3a-driven GFP delineates Gustatory Fibers innervating Sour-responsive Taste Cells: A Labeled Line for Sour Taste? J Comp Neurol. 525:2358–2375.

Stuart, T., Butler, A., Hoffman, P., Hafemeister, C., Papalexi, E., Mauck, W.M., Hao, Y., Stoeckius, M., Smibert, P., and Satija, R. 2019. Comprehensive Integration of Single-Cell Data. Cell. 177:1888–1902.e21.

Taruno, A., Vingtdeux, V., Ohmoto, M., Ma, Z., Dvoryanchikov, G., Li, A., Adrien, L., Zhao, H., Leung, S., Abernethy, M., et al. 2013. CALHM1 ion channel mediates purinergic neurotransmission of sweet, bitter and umami tastes. Nature. 495:223–226.

Vandenbeuch, A., Larson, E.D., Anderson, C.B., Smith, S.A., Ford, A.P., Finger, T.E., and Kinnamon, S.C. 2015. Postsynaptic P2X3-containing receptors in gustatory nerve fibres mediate responses to all taste qualities in mice. J Physiol. 593:1113–1125.

Zerbino, D.R., Achuthan, P., Akanni, W., Amode, M.R., Barrell, D., Bhai, J., Billis, K., Cummins, C., Gall, A., Girón, C.G., et al. 2018. Ensembl 2018. Nucleic Acids Res. 46:D754–D761.

Zhang, J., Jin, H., Zhang, W., Ding, C., O’Keeffe, S., Ye, M., and Zuker, C.S. 2019. Sour Sensing from the Tongue to the Brain. Cell.

